# Water availability determines plant regeneration fates

**DOI:** 10.1101/2024.07.30.605771

**Authors:** Abdul Kareem, Anna K. van Wüllen, Ai Zhang, Gabriel Walckiers, Ellen Fasth, Charles W. Melnyk

## Abstract

Wounding and hormones serve as diverse triggers for regeneration in animals and plants but how organisms determine regeneration outcomes remains largely unknown. Here, we demonstrated that wounded Arabidopsis tissues regenerate two distinct fates, wound-induced callus or de novo root formation, that are driven by antagonizing molecular pathways related to cambium and root development. We discovered that local water availability dictated these regeneration outcomes in Arabidopsis and tomato, with high water triggering root fate and low water initiating callus fate. Moreover, distinct spatial distributions of auxin response maxima were critical for fate progression and water availability regulated these auxin maxima through the hormones ethylene and jasmonic acid. We propose that water availability determines environmental control of regeneration plasticity with applied potential for improving regeneration in agriculture.

## Main text

Plants adapt to fluctuating environmental conditions, a remarkable trait largely controlled by their developmental plasticity. Such plasticity relies on cell fate determination by internal hormonal signals, transcription factors and receptors as well as the external environmental factors such as water availability, nutrient supply and temperature. For instance, roots employ hydropatterning to form lateral roots in response to local water availability (*3-5*). Similarly, a population of pericycle cells within roots display high plasticity to give rise to lateral roots, cambium or periderm depending on root developmental age and hormonal cues (*6-8*). Such pericycle-derived cells express transcription factors such as *LB?>D16* when obtaining lateral root fates (*7, 9*), whereas they express *LBD11* and *WOX4* when obtaining cambium fates (*6, 7, 10, 11*).

This developmental plasticity also underpins the regenerative capacity of plants, crucial for biotechnology and plant propagation. Culturing plant tissues on hormone-rich medium induces pericycle cells to form undifferentiated cell masses called callus which, by changing the ratio of auxin and cytokinin, can be subsequently regenerated into tissues such as shoots or roots (*12-14*). Such hormone induced callus (HIC) expresses root stem cell regulators *PLT1* and *PLT2* and occurs via molecular pathways resembling root meristem development (*15, 16*). However, when plant tissues are wounded, they also trigger a regeneration process, for example, wound-induced callus formation (WIC), *de novo* root regeneration (DNRR), root tip healing or vascular reconnection (*17-22*). Such regeneration pathways allow plants to recover from injury without external hormonal requirement. With WIC, activation of transcription factors like *WIND1* and *WOX13*, along with cytokinin, initiates callus formation which seals the wound and later differentiates to missing tissues (*19, 23, 24*). During DNRR, auxin activates *WOX11* and *LBD16* transcription factors to facilitate adventitious root formation from wounded tissues (*18, 25, 26*). Although the molecular players involved in different regeneration processes are well known, how plants determine whether to trigger a specific regeneration fate remains unknown. Moreover, unlike DNRR and HIC, the molecular fate of WIC remains unresolved.

Here, by investigating diverse regeneration fates initiated from the same tissue, we demonstrated a trade-off between different forms of regeneration driven by distinct cambium or root related molecular pathways. We discovered that water availability is a factor deciding whether plant tissues undergo root-mediated or callus-mediate regeneration fates in both Arabidopsis and tomato. Furthermore, we revealed the spatial distribution of auxin response maxima is relevant for such fate changes and that water availability regulates such auxin maxima through the hormones ethylene and jasmonic acid.

## Results

### Different regeneration fates use different molecular pathways

The same plant tissue can exhibit different types of regeneration, so we sought to determine whether different regeneration fates had common or distinct molecular pathways. We compared three different regeneration fates from Arabidopsis leaf petioles: wound-induced callus formation (WIC), wound-induced *de novo* root regeneration (DNRR), and unwounded hormone-induced callus formation (HIC). DNRR was initiated at the wound site of the petiole touching hormone-free culture medium, while WIC was induced at the wound site when not in contact with the medium (Fig. 1A-D), similar with previous findings (*17*). HIC, on the other hand, was triggered by culturing unwounded tissues in an auxin-rich medium (Fig. 1E). We focused on key genes implicated in DNRR and observed activation of *LBD16, PLT2*, and *WOX11* during both DNRR and HIC (Fig. 1B,C and E and fig. S1A,B), while their expression was low in WIC (Fig. 1D and fig. S1C). We monitored the expression patterns of genes associated with the vascular cambium and found *LBD1, LBD11, PXY*, and *WOX4* activated in the proliferating cells of WIC (Fig. 1D and fig. S1C), while their expression levels were lower during both DNRR and HIC (Fig. 1C and E and fig. S1B). The similar expression profiles of DNRR and HIC suggest that they likely shared similar molecular pathways, while WIC likely used a pathway linked to the vascular cambium.

**Fig. 1:**
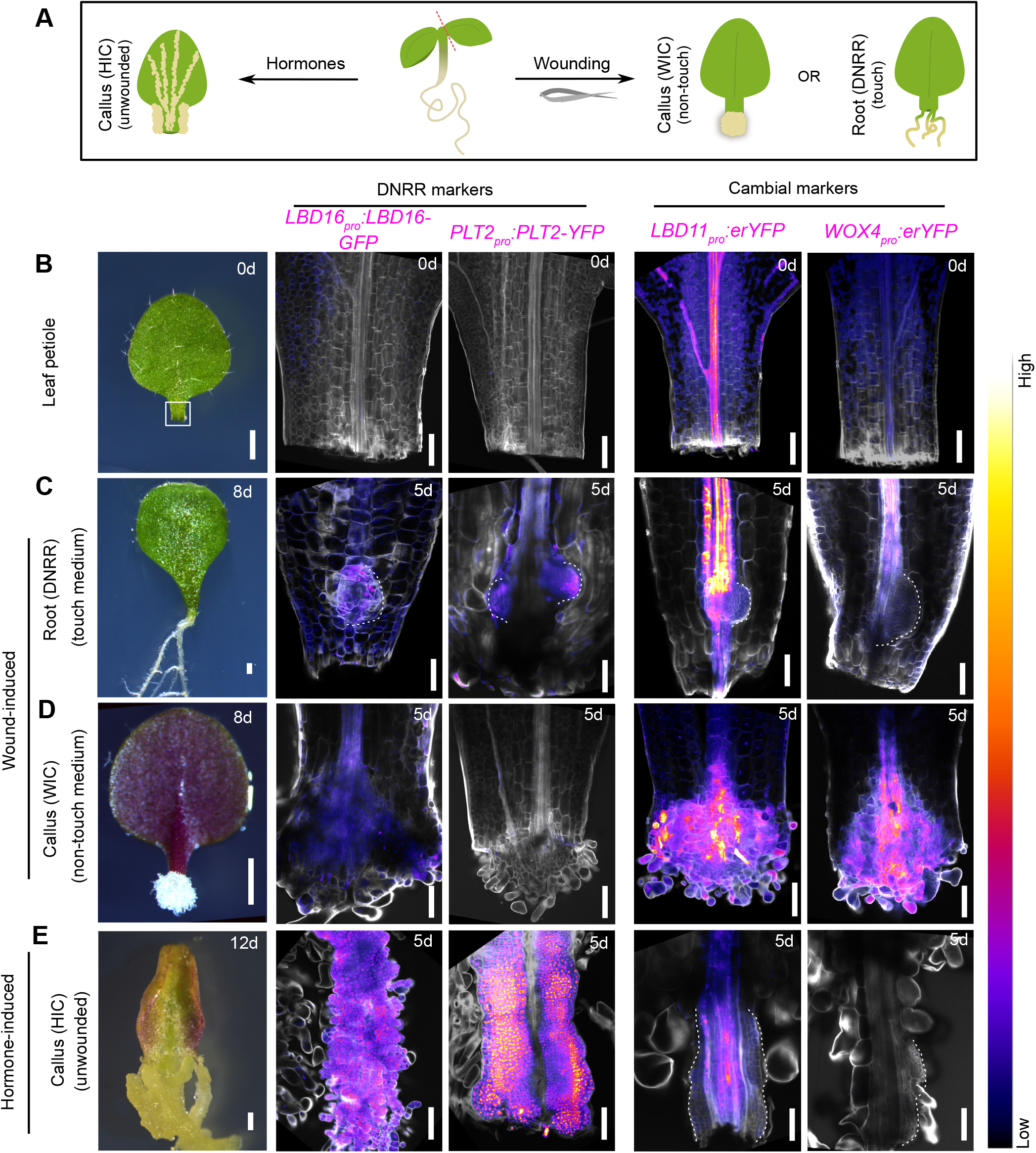
Different regeneration fates use different molecular pathways. (**A**) Schematic showing diverse plant regeneration modes: hormone-induced callus formation (HIC), wound-induced callus formation (WIC) and wound-induced *de novo* root regeneration (DNRR). (**B-E**) Expression patterns of fluorescence reporters (magenta) for *LBD16, PLT2* (DNRR markers), *LBD11* and *WOX4* (cambial markers) in the Arabidopsis leaf petiole on day 0 (B) or during DNRR (C), WIC (D), or HIC from unwounded tissues (E) on day 5. Dashed lines indicate the regenerating root primordium in (C) and callus in (E). Calcofluor white (grey) was used to stain cell walls. The corresponding representative bright-field images of the detached leaf (B), DNRR (C), WIC (D) and HIC (E) at 0, 8 or 12 days (d) are shown in the left panel. Scale bar: 500µm for the bright-field images, 100µm for the confocal images.

### Wound-induced callus inhibits root regeneration

To further investigate the molecular basis for WIC, we tested cambial mutants, including *lbd1,11*; *lbd1,3,4,11*; and *wox4,14*, and found they were defective in WIC formation (Fig. 2A and B and fig. S2A), suggesting an essential role of cambial genes in this process. Inducible overexpression of *LBD11* or *WOX4* also boosted WIC formation (Fig. 2B and fig. S2B). Notably, the pericycle cell division-defective mutant *solitary root1* (*slr-1*) did not inhibit WIC formation (fig. S2C). Despite deficient WIC formation, cambial mutants displayed an enhanced ability to form DNRR, even in conditions where wild-type tissues failed to form roots (Fig. 2A and C and fig. S2A and D-F). We observed increased DNRR-related gene transcription in the WIC deficient *lbd1,11* mutant (fig. S2G), implying suppression of WIC activated DNRR-related pathways. Additionally, overexpression of WIC-promoting genes *LBD11* and *WOX4* inhibited the DNRR pathway, while overexpression of DNRR-promoting genes *LBD16, PLT2*, and *WOX11* suppressed the WIC pathway (Fig. 2D and E and fig. S2H and I). Our data indicate that activation of cambial cells were crucial for WIC formation and that WIC fate inhibited DNRR fate, suggesting a trade-off between regeneration fates.

**Fig. 2:**
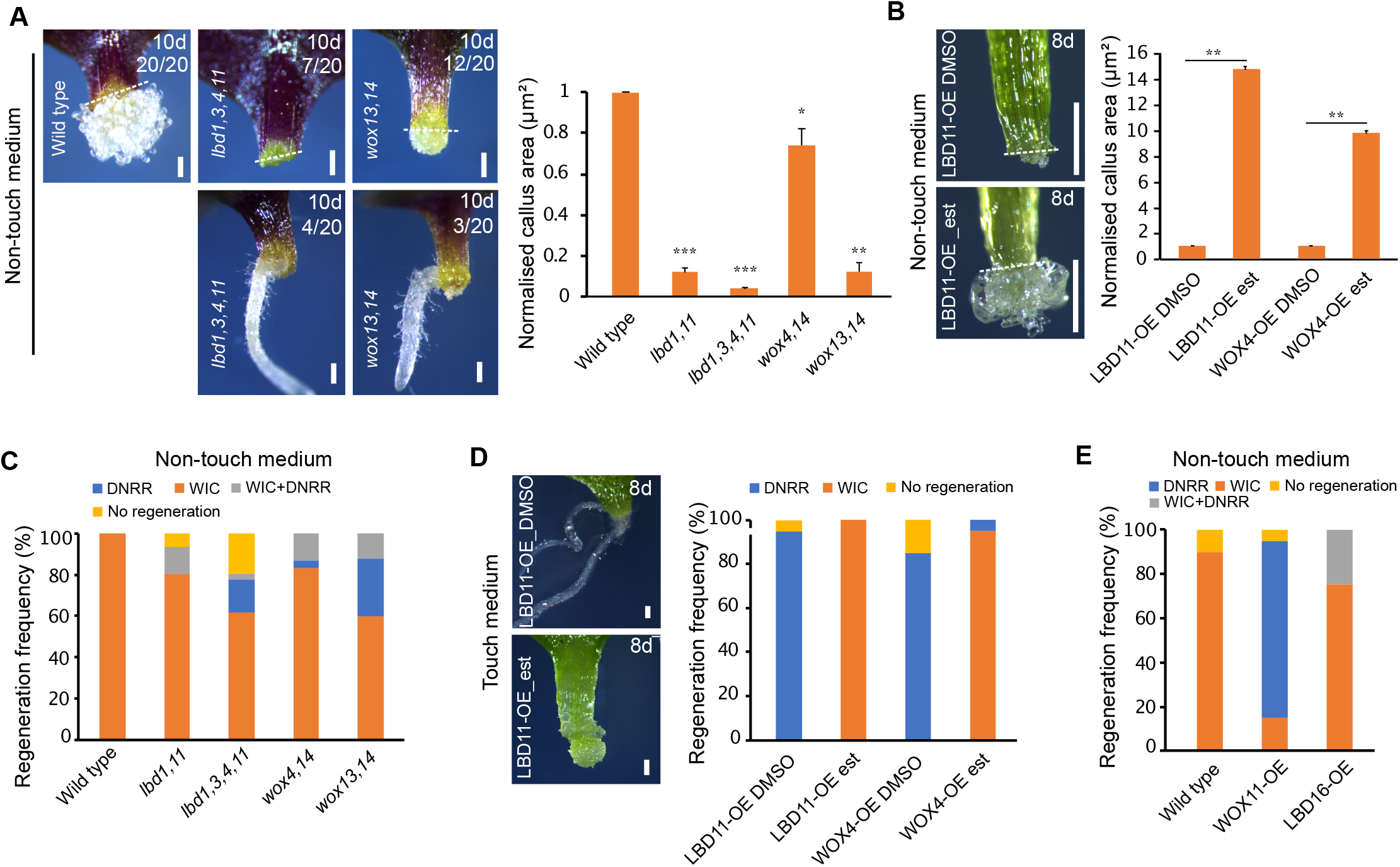
Wound-induced callus inhibits root regeneration. (**A**) Top panels showing wound-induced callus (WIC) formation in leaf petiole explants of Arabidopsis wild type and cambial mutants, with the wound site not touching the medium. Bottom panels showing *de novo* root regeneration (DNRR) in the mutants under the same conditions. Numbers on the image indicate how many plants out of 20 showed the phenotype. Normalised WIC area quantifications of wild type and mutant explants are shown to the right (n=20-24 explants per genotype; mean±SD; *p<0.05; **p<0.01; ***p<0.001; Student’s t-test). (**B**) WIC formation images or quantifications in the hypocotyl explants of *LBD11-OE* or *WOX4-OE* with the wound site not touching the medium. Mock (DMSO) and 5µM estradiol (est) treatments are shown (n=10 explants per treatment per genotype; mean±SD; **p<0.01; Student’s t-test). (**C**) Frequency of regeneration of WIC and DNRR in the leaf petioles of various mutants with the wound site not in contact with the medium (n=21-25 explants per genotype). (**D**) WIC formation images or quantifications in the leaf explants of *LBD11-OE* or *WOX4-OE* when the wound site is in contact with the medium. Mock (DMSO) and 5µM estradiol (est) treatments are shown (n=20 explants per genotype; mean±SD; Student’s t-test). (**E**) Regeneration frequencies of DNRR and WIC in the leaf petiole of *LBD16-OE* (constitutive) and *WOX11-OE* (constitutive) with the wound site not touching the medium. (n=20 explants per genotype; mean±SD; Student’s t-test). Dashed lines indicate the wound site in (A and B). Scale bar: 200µm

### Trade-off between regeneration of callus and root is determined by water availability at the wound site

What determined cell fate was unknown but we observed that when the wound site was in contact with B5 medium, DNRR was triggered (Fig. 3A and C), while contact with parafilm or air favored WIC formation (Fig. 3A and C) suggesting that physical contact alone did not influence regeneration fates. Nutrient availability also had no significant impact (fig. S3A and B). We then tested the role of water by culturing leaf tissues with the wound site in contact with B5 medium with various agar concentrations (0.375% to 3%) which represented different water potentials (fig. S3C). This variation corresponded to the available water at the medium surface, as previously described (*5*). We found higher water availability (lower agar concentrations) favored DNRR, while lower water availability (higher agar concentrations) promoted WIC (Fig. 3A and C). Applying water to the wound site touching parafilm induced DNRR, whereas parafilm alone led to WIC formation (Fig. 3A and C). This fate decision was made shortly after wounding, and changes in water availability during this period impacted regeneration outcomes (fig. S3D). Similar to Arabidopsis, tomato tissues also responded to water availability by regenerating roots or callus (Fig. 3B and D). Air layering is commonly used to propagate plants, so we used this technique on tomato stems and found damp soil promoted DNRR whereas dry soil promoted WIC (Fig. 3E and F) supporting the observation that water availability, rather than physical contact, controlled regeneration outcomes. To further understand the genes involved in fate decision, we conducted RNA-sequencing (RNA-seq) from wounded Arabidopsis petioles at various time points with two agar concentrations (Fig. 3G and fig. S4). DNRR-promoting genes (*LBD16, PLT2, WOX5*) were induced under high water conditions, while WIC-promoting genes (*LBD11, WOX4, WOX14*) were upregulated under low water conditions (Fig. 3H, fig. S3E). The corresponding mutants displayed defects in either WIC or DNRR, consistent with a role for promoting either DNRR or WIC during varying water availability (fig. S3F and G). Overall, our findings suggest that water availability at the wound site was crucial for deciding whether tissues undergo root or callus regeneration in Arabidopsis and tomato.

**Fig. 3:**
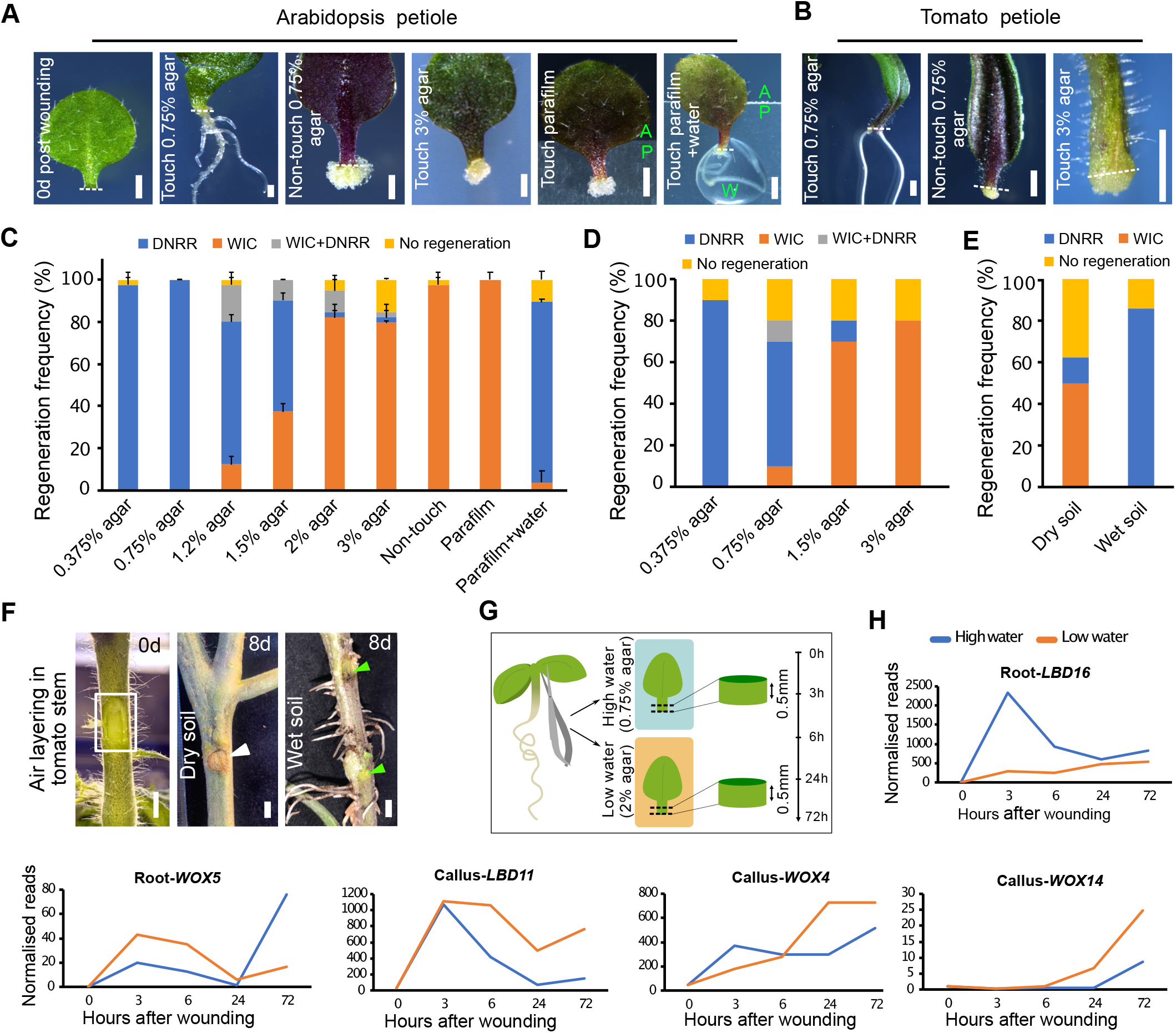
The trade-off between regeneration of callus and root is determined by the availability of water. (**A**) Regeneration in Arabidopsis leaf petiole tissues with varying agar concentrations or medium contact, 10 days (d) post-injury. A 0 day control is included as well as parafilm contact and moistened parafilm contact. Dashed line marks the wound site. ‘A’ = agar, ‘P’ = parafilm, ‘W’ = water. (**B**) Regeneration in tomato cotyledon petiole tissues with varying agar concentrations or medium contact. Dashed line indicates the wounded site. (**C** and **D**) Frequency of regeneration in Arabidopsis leaf petiole (C) and tomato cotyledon petiole (D) demonstrating either *de novo* root regeneration or wound-induced callus under different water availability conditions. (n=20 explants per genotype; mean±SD; Student’s t-test). (**E** and **F**) Frequency of regeneration and images of tomato stems during air-layering with either wet or dry soil. (n=20 explants per genotype; mean±SD; Student’s t-test). Wound sites are indicated with a rectangle box in the 0d image and a green arrowhead in the 8d wet soil image. Callus formation is indicated by a white arrow head. (**G**) Schematic depicting 0.5mm tissue harvesting and time points for Arabidopsis petiole wound sites under high and low water conditions for RNA-seq. (**H**) Transcriptional dynamics of genes associated with callus and root regeneration in Arabidopsis. Scale bar: 500µm in (A), 1mm in (B) and 5mm in (F).

### Water availability shapes spatial auxin distribution during regeneration

Our transcriptome data revealed differential expression of many auxin-responsive genes under varying water availability (fig. S5A). We therefore explored auxin distribution using the auxin responsive reporter DR5-VENUS and observed distinct spatial expression patterns in Arabidopsis tissues across water conditions. High water availability (0.75% agar) caused the auxin maxima to form at the root primordium initiation site, distal to the wound, whereas low water availability (2% agar) caused it to form near the wound site (Fig. 4A and fig. S5B). Intermediate water levels (1.5% agar), promoting both DNRR and WIC regeneration, resulted in a broader auxin distribution (Fig. 4A).

**Fig. 4:**
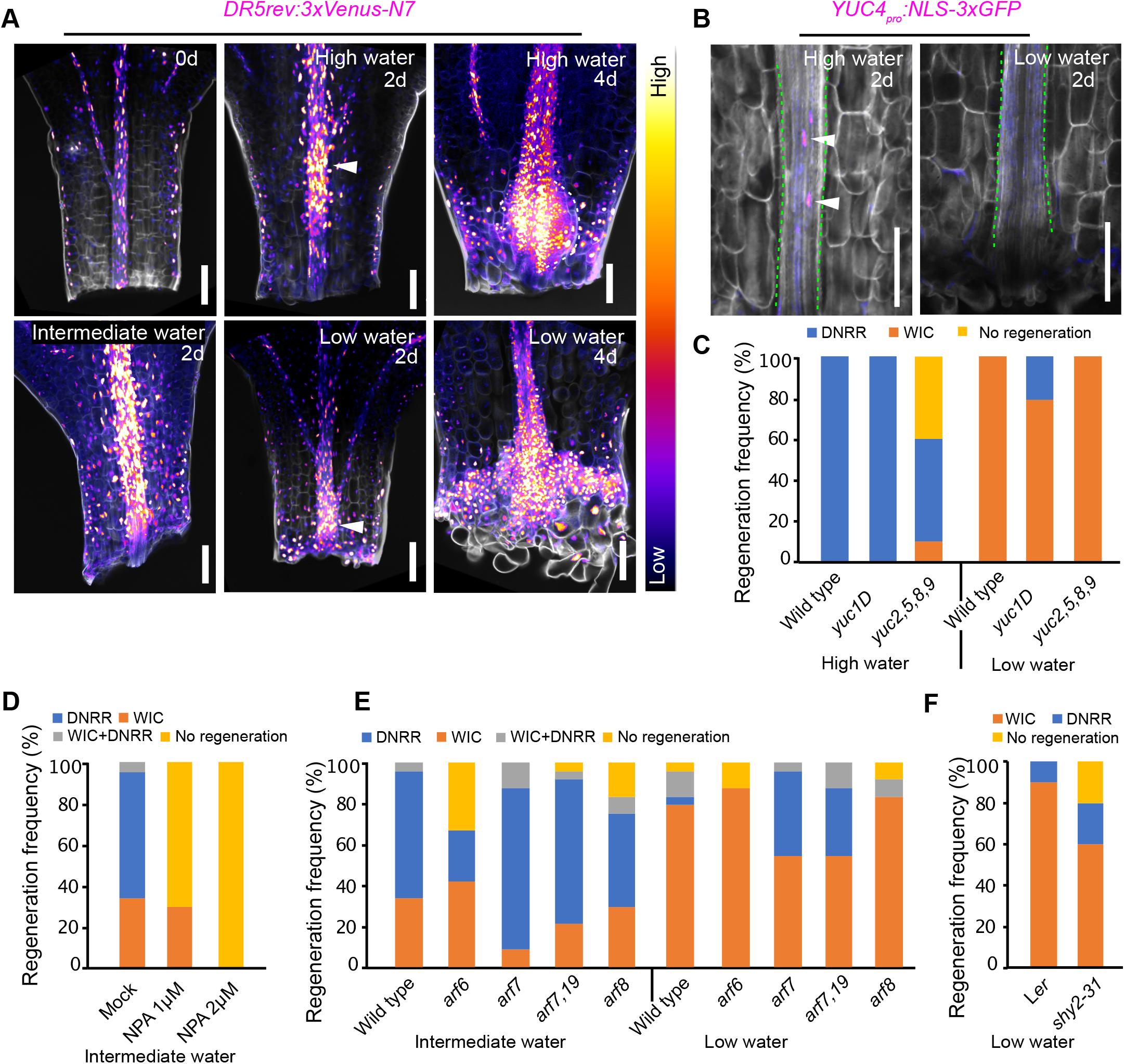
Water availability shapes spatial auxin response during regeneration. (**A**) *DR5-VENUS* auxin responsive reporter under high (0.75% agar), intermediate (1.5% agar) and low (2% agar) water conditions, with arrowheads indicating auxin maxima in the Arabidopsis leaf petiole. (**B**) Expression of *YUC4-GFP* under high (0.75% agar) and low (2% agar) water conditions. Green dotted lines mark the vasculature. (**C**) Regeneration frequencies in YUCCA1 over expressor (*yuc1D*) and *yuc2,5,8,9* mutants at high and low water conditions (n=19-20 explants per genotype; mean±SD; student’s t-test). (**D**) Regeneration frequencies after NPA treatment (1µM and 2µM) at intermediate (1.5% agar) water conditions (n=10-16 explants per treatment). (**E**) Regeneration frequencies in *arf6, arf7* and *arf8* mutants at intermediate (1.5% agar) and low (2% agar) water conditions (n=24 explants per genotype). (**F**) Regeneration frequencies in the *shy2-31* mutant under low water conditions (3% agar) (n=10 explants per genotype). Scale bar= 100µm.

Auxin biosynthesis genes such as *ASA1, YUC1, YUC4*, and *YUC5* were upregulated at high water, with *YUC4-GFP* activating specifically at the root primordium initiation site (Fig. 4B and fig. S5C). We tested a role for auxin biosynthesis and found the *yuc2,5,8,9* mutant impaired DNRR but had normal WIC formation. Treatment with exogenous auxin or using the auxin overproducing line *yuc1D* perturbed WIC but enhanced DNRR (Fig. 4C and fig. S5D-F). However, treatment with the auxin transport inhibitor NPA revealed that auxin transport was crucial for both processes, with DNRR being more sensitive to its perturbation (Fig. 4D). We investigated downstream auxin signaling and found mutants *arf6* and *arf8* displayed defects in DNRR, while *arf7* and *arf7,19* mutants showed an increased ratio of DNRR to WIC formation (Fig. 4E). SHY2-VENUS, a reporter of a negative auxin regulator, displayed distinct spatial distribution, with low signals at high water and high signals at low water (fig. S5G). The loss-of-function *shy2-31* mutant promoted DNRR under low water, indicating that high SHY2 activity inhibits DNRR (Fig. 4F). Thus, there appeared to be pathways involving DNRR-promoting ARFs and WIC-promoting ARFs. Taken together, water availability dictated regeneration fates through auxin biosynthesis, transport, and signaling pathways which likely influenced spatial auxin distribution.

### Water availability regulates stress hormones to influence spatial auxin distribution and regeneration

Stress hormones are important for early wound response (*27, 28*) and we observed activation of genes responsive to ethylene and jasmonate in our RNAseq datasets within 3 hours of wounding under high water conditions (Fig. 5A, fig. S6A). Reporter gene analyses also showed upregulation of ethylene biosynthesis (ACS6-VENUS) and jasmonate response (JAZ10-VENUS) markers in high water conditions (Fig. 5B). Consistently, short treatments of ethylene (4µM ACC) and methyl-jasmonate (5µM MeJA) enhanced the DNRR to WIC ratio (Fig. 5C). Additionally, treatment with the ethylene inhibitor silver nitrate suppressed DNRR and promoted WIC formation ratio (Fig. 5D). This effect was partially rescued by auxin (NAA) treatment (Fig. 5D).

**Fig. 5:**
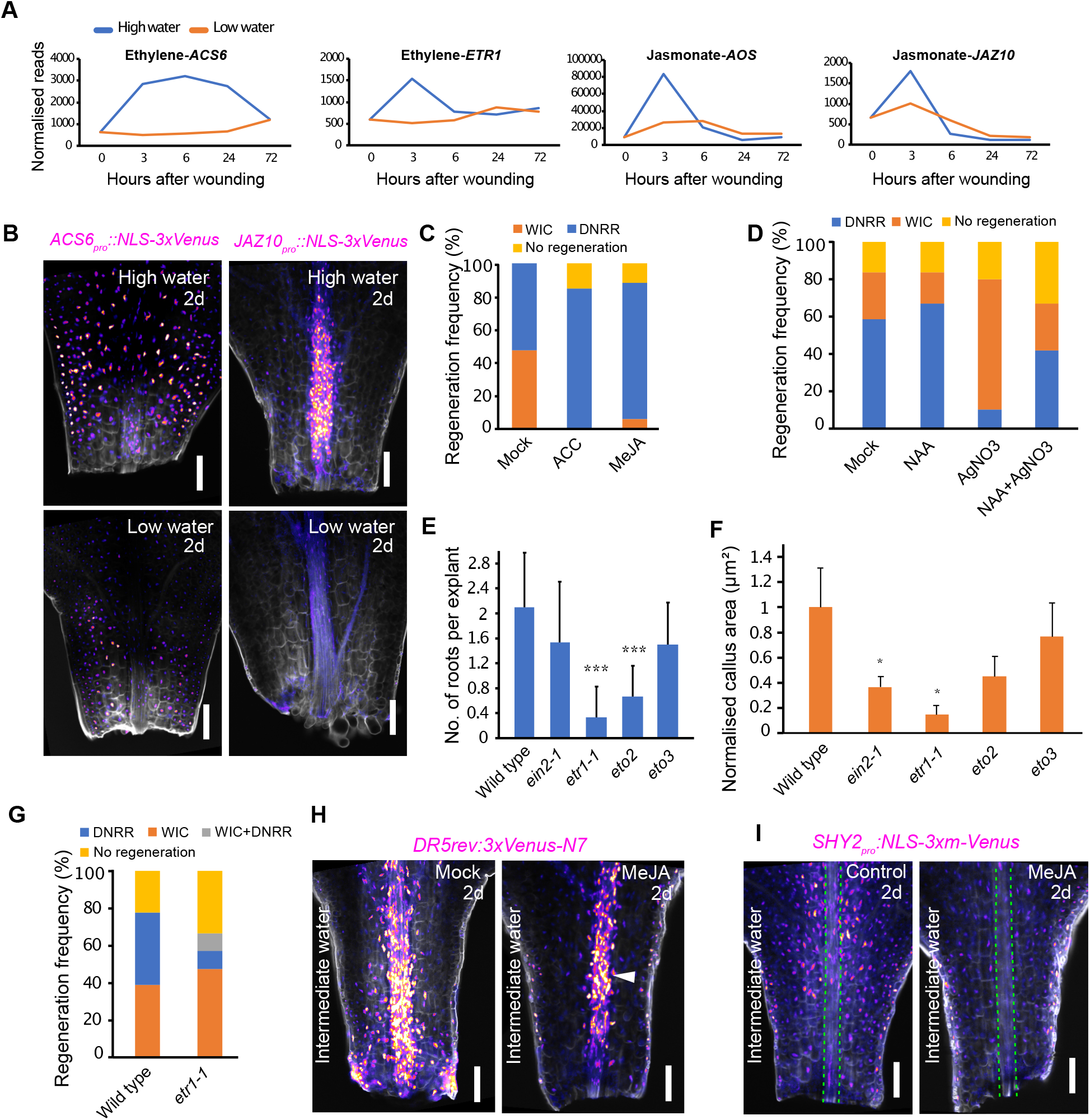
Water availability regulates stress hormones to influence spatial auxin distribution. (**A**) Transcriptional dynamics of genes associated with ethylene and jasmonate synthesis and signalling. (**B**) Ethylene biosynthesis reporter *ACS6-VENUS* and jasmonate-response reporter *JAZ10-VENUS* expression under high (0.75% agar) and low (2% agar) water conditions. (**C**) Regeneration frequencies after treatment with ethylene (4µM ACC) or methyl-jasmonate (5µM MeJA) at intermediate water conditions (1.5% agar). (n=17-20 explants per treatment). (**D**) Regeneration frequencies after treatments with 0.2µM NAA, 10µM AgNO3 or NAA+AgNO3 at intermediate water conditions. (n=10-18 explants per treatment). (**E**) Frequency of root regeneration in ethylene mutants compared to wild type under high water conditions (n=10-13 explants per genotype; mean±SD; ***p<0.001; Student’s t-test). (**F**) Normalised wound-induced callus (WIC) area of wild type and ethylene mutants under low water conditions (n=6-11 explants per genotype; mean±SD; *p<0.05; Student’s t-test). (**G**) Regeneration frequency in the *etr1-1* mutant under intermediate water conditions. (n=18-21 explants per genotype). (**H**) *DR5-VENUS* expression after mock or 5µM MeJA treatment for 43h at intermediate water conditions (1.5% agar). (**I**) *SHY2-VENUS* expression after mock or 5µM MeJA treatment for 43h at intermediate water conditions (1.5% agar). Green dotted lines mark vasculature. Scale bar: 100µm.

The ethylene signaling mutant *ethylene resistant1-1* (*etr1-1*) (*29*) exhibited defects in both DNRR and WIC (Fig. 5E,F). However, it displayed a greater impairment in the DNRR fate, leading to a higher WIC to DNRR ratio at intermediate water availability (Fig. 5G). Notably, the ethylene overproducing mutant *eto2* (*30*) also affected DNRR, with minimal impact on WIC (Fig. 5E,F). So although our transcriptome and treatment assays indicated a strong involvement of ethylene and jasmonate in DNRR, some ethylene signaling appeared relevant for both DNRR and WIC.

Using the DR5-VENUS reporter, we observed that ethylene and jasmonate directly influenced spatial auxin distribution. ACC or MeJA treatments resulted in more confined auxin maxima, mimicking high water conditions under intermediate water availability (1.5% agar), compared to the typically broader distribution normally observed (Fig. 5H and fig. S6B). Furthermore, MeJA treatment reduced SHY2-VENUS signal at intermediate water availability, similar to high water conditions that promote DNRR fate (fig. S6C). Our findings suggest that stress hormones ethylene and jasmonate, activated soon after wounding in response to water availability, influenced spatial auxin distribution and promoted DNRR regeneration fates (fig. S6D).

## Discussion

When tissues detach from a growing plant, they often lose access to water, nutrients, growth signals, and positional cues, necessitating their reliance on internal and external stimuli for survival and adaptation. Our study reveals external water availability as a key factor determining regeneration fates in wounded tissues, overshadowing in our study the influence of nutrients and physical contact observed in other developmental processes (*31-33*). Mechanical pressure along with developmental age and positional information all impact cell fate changes during restorative regeneration, (*34, 35*). Physical contact may also influence regeneration (*17*), however, our study found that surface contact alone was inadequate to determine regeneration fates in detached tissues. This could be due to detached tissues or tissues with large wounds facing a restricted environment from lost connections, missing positional information, and increased dehydration unlike intact tissues undergoing restorative regeneration (*34, 35*). Consequently, detached tissues rely on water signals, important for sustaining cell viability, to determine their regeneration fates leading to regeneration of either root for water uptake or pluripotent callus for wound sealing. Severe damage across multiple cell layers in mature tissues could also manifest detached tissue behavior to fate determination, as demonstrated in air layering. Thus, water signals appear to exert greater influence over other signals on detached and severely damaged tissues.

Water availability is also crucial for root branching in lateral root hydropatterning (*5*). Water dictates a single developmental fate in root hydropatterning, and shows irreversible changes with short term deficiency (*3*). In contrast, here we show that water determines a trade-off between two regeneration fates operating through distinct pathways, offering some reversibility. This underscores the greater flexibility of our findings with regeneration. Furthermore, regenerating tissues respond to fluctuating water availability rather than mere presence or absence, contrasting with xerobranching, an extreme form of hydropatterning, wherein water absence represses root branching (*5, 36*). We found this narrow window of water levels influenced the distinct spatial distribution of auxin response maxima in regeneration pathways by coordinating with stress hormones such as ethylene and jasmonate. We uncovered that transient elevations in these stress hormones were crucial for regeneration, consistent with previous work (*27, 28*), but prolonged activity inhibited both regeneration fates, likely triggering defense pathways (*37*). This suggests a potential trade-off between regeneration and defense (*38*), where the initial surge in stress hormones promotes regeneration, yet prolonged activation may trigger defensive responses. Such defense responses could be an extreme manifestation of adaptive responses. However, further investigation is required to understand this correlation and explore whether extreme water levels or water deprivation also trigger a defense-like response, potentially linking to mechanisms like xerobranching. Altogether, our data provide mechanistic insight into water availability-dependent regeneration pathways through hormonal crosstalk, opening new research avenues.

Previously, it was shown that liquid or low agar medium enhances shoot and root organogenesis in tissue culture for several species, including *Picea abies* and Sweetgum (*39, 40*). Our findings provide a mechanistic basis for this enhanced tissue culture method and reveal new possibilities for enhancing regeneration by modifying water availability. Additionally, fine-tuning stress hormones such as ABA, ethylene, and jasmonate through transient exposure or low concentrations holds promise for improving regeneration, particularly in somatic embryogenesis where use of stress hormones is common (*41, 42*) and also with wound-induced propagation. Thus, our research paves the way for significant advancements in agricultural regeneration techniques.

## Materials and Methods

### Plant materials and growth conditions

*Arabidopsis thaliana* ecotype Columbia-0 (Col-0) was used throughout as the wild type, unless otherwise indicated. Tomato (*Solanum lycopersicum*) variety Moneymaker was used. All Arabidopsis mutants and transgenic lines used in this study have been previously described and the details are listed in Table S1. Arabidopsis seeds were surface-sterilized with 70% ethanol for 10 minutes and dried on sterilized filter paper. The seeds were then plated on half-strength Murashige and Skoog (MS) medium with 0.5% sucrose, 0.25 g l^−1^ MES buffer pH 5.7–5.75 and 0.8% (w/v) plant agar unless otherwise indicated. After 2 days of stratification at 4 °C in the dark, seedlings were grown vertically at 22 °C under a 16-h light (110μE intensity) and 8-h dark photoperiod.

Tomato seeds were surface sterilized with 75% bleach for 10-15 minutes, washed five times in water, and plated on MS medium containing 0.5% sucrose, 0.25 g/l MES buffer (pH 5.7–5.75), and 0.8% (w/v) plant agar unless otherwise indicated. The plates were then transferred to a growth cabinet at 25 °C with a 16-hour light and 8-hour dark photoperiod.

### Wounding and regeneration assays

Cotyledon, leaf, and hypocotyl explants were employed for wound-induced regeneration in both Arabidopsis and tomato. Cotyledons with approximately 0.5 mm petioles were carefully detached from 7-day-old seedlings using micro scissors. Hypocotyls were wounded above the root-hypocotyl junction in 7-day-old seedlings. For leaf wounding, the first pair of leaves with petioles (0.5 mm) were detached from 10-day-old seedlings. The detached explants were subsequently transferred to hormone-free Gamborg B5 medium containing 2% glucose, 0.5 g l–1 MES buffer (pH 5.7), and 0.75% (w/v) plant agar. The explants were oriented with their wound site either touching the medium (abaxial side facing the medium for leaf and cotyledon) or not touching the medium (adaxial side facing the medium for leaf and cotyledon), following the approach described previously (*17*). In the not-touching condition, the leaf and cotyledon explants were also oriented with the abaxial side after removing an agar block from the medium near the wound site, aiming to test the specific effect of wound-site orientation over leaf side orientation (abaxial or adaxial). The explants were cultured horizontally for regeneration under the same growth conditions as mentioned above. Regeneration responses were analyzed between 6 days and 10 days after wounding. For the regeneration response on parafilm, the detached leaf explants were oriented with their wound site in contact with a parafilm strip placed on B5 medium. For regeneration response to water availability, the detached leaf explants were transferred onto the B5 medium with varying concentrations of agar (0.375%, 0.76%, 1.5%, 2% and 3% plant agar (Duchefa Biochemie)). The water potential of the agar plates was calculated using the following formula after measuring the water activity (aw) and temperature (T) of the different agar media with a water activity meter (AquaLab Pre)

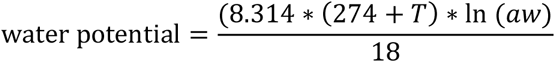

Water potential measurements were made in triplicate for each agar concentration and averaged.

For regeneration on nutrient-free medium, the detached leaves were cultured on a medium containing only agar without added nutrients. For regeneration on hormone medium, ten-day-old whole seedlings were transferred onto a B5 medium supplemented with 2 μg/ml of 2,4-dichlorophenoxyacetic acid (2,4-D) and 0.05 μg/ml of kinetin for hormone-induced callus induction (HIC), without any wounding. After five days of culture, the petiole calli were harvested for confocal imaging. For air-layering in tomato, the stem was wounded using a sharp blade in the greenhouse. The wounded area was then wrapped with plastic film containing 5g of either dry or wet soil, and tissue regeneration was monitored after two weeks.

### Chemical treatments

For the inducible overexpression of *LBD11* and *WOX4, 35S::XVE>>LBD11* and *35S::XVE>>WOX4* seedlings were pre-treated with B5 medium containing 5 µM estradiol for 6 hours before wounding. After wounding, tissues were placed on the same medium, either touching or not touching it. An equivalent volume of DMSO was added to the B5 medium for the mock treatment. For *PLT2* overexpression, *35S::PLT2-GR* petiole tissues were cultured on B5 medium with 5 µM dexamethasone (DEX) with the wound site not touching the medium. Ethanol was used for the mock treatment. For auxin treatments, petiole tissues were cultured on B5 medium with 0.2 µM 1-naphthaleneacetic acid (NAA) or DMSO (mock) under high (0.75% agar) or intermediate (1.5% agar) water conditions. For inhibition of polar auxin transport, 1 µM or 2 µM N-1-naphthylphthalamic acid (NPA) or DMSO (mock) was added to the B5 medium and cultured the petiole tissues under high or intermediate water conditions. For ethylene and jasmonic acid treatments, 4 µM 1-aminocyclopropane-1-carboxylate (ACC) or 5 µM methyl-jasmonate (MeJA) was added to the B5 medium, with water or ethanol, respectively, as the mock. Petiole explants were cultured for 24 or 43 hours under high or intermediate water conditions and then transferred onto hormone-free B5 medium under the same water conditions. For inhibition of ethylene activity, 10 µM silver nitrate (AgNO3) or water (mock) was added to the B5 medium and cultured the petiole tissues under high or intermediate water conditions. For the NAA+AgNO3 treatment, 0.2 µM NAA and 10 µM AgNO3 were added together to the medium, and the petiole tissues were cultured under intermediate water conditions.

### Tissue fixation and clearing for confocal imaging

The regenerating tissues with various fluorophores were fixed, cleared and then stained for cell walls before confocal imaging using the previously described protocol(*43*) with modifications. Briefly, tissues were fixed with 4% PFA in 1X PBS buffer with 0.1% Triton-X 100 for 1h at room temperature, including a 10-minute vacuum application. After fixation, the tissues were washed twice in 1X PBS, followed by treatment with absolute methanol and then absolute ethanol for 30 minutes each. Subsequently, after a wash with 1X PBS, the tissues underwent clearing in ClearSee solution (10% xylitol, 15% sodium deoxycholate, 25% urea) at 4 °C for 2-3 days. The tissues were then stained in 0.1% Calcofluor white in ClearSee solution for 30-60 minutes, followed by destaining in ClearSee for 15 minutes.

### Microscopy

Confocal images were captured using a Zeiss LSM780 with a 20X dry objective (N.A 0.8). The pixel format was set to either 512×512 or 1024×1024 (for high-resolution images). A bidirectional scan was configured with a scan speed of 8, employing line averaging for 4. The following wavelength settings were used for imaging various fluorophores. For calcofluor white imaging, excitation wavelength (ex.) 405 nm; emission (em.) 414-471 nm. For CFP, ex. 458 nm; em. 464–499 nm; For GFP: ex. 488 nm; em. 491–519 nm. For YFP and Venus: ex. 514 nm; em. 518–540 nm. For RFP and propidium iodide: ex. 561 nm; em. 585–630 nm. The laser power and pinhole were adjusted based on fluorescence brightness to prevent signal saturation or bleaching. The gain was set to 1. Imaging of CFP/GFP/YFP/RFP was conducted using ChS1 detector. The settings remained unchanged throughout each experiment where fluorescent signal comparison was needed.. Bright-field images of regenerating callus and *de novo* roots were captured with a Leica M205FA stereo microscope.

### Image analysis and data processing

The images were analyzed using FIJI software (https://fiji.sc). The callus area was measured by the freehand tool in FIJI. Images were sometimes rotated for better orientation. The brightness of the fluorescence signal was adjusted for improved visualization, and these adjustments were consistently equal across various time points or treatments. Images were then annotated and arranged in Affinity Designer (version 1.9.1).

### RT-qPCR assays

Arabidopsis leaf petioles from wild-type and *lbd1,11* mutant plants were wounded and cultured for regeneration with the wound site not in contact with the medium. Approximately 0.5 mm of tissue from the wound site was collected 4 days after wounding. Total RNA was extracted using a Roti-Prep RNA MINI Kit, and the RNA samples were quantified with a NanoDrop ND-1000 spectrometer (Thermo Fisher Scientific). For cDNA synthesis, 500 ng of total RNA was reverse-transcribed using the Maxima First Strand cDNA Synthesis Kit, which includes oligo(dT) and random hexamer primers. Quantitative PCR (qPCR) was carried out with the iCycler iQ Real-Time PCR detection system in 10 μL reaction volumes. Each reaction contained 5 μL of 2X Maxima SYBR Green qPCR/ROX Master Mix, 0.75 μM of both forward and reverse primers, and 2 μL of diluted cDNA. The qRT-PCR protocol included an initial denaturation at 95°C for 10 minutes, followed by 40 cycles of 95°C for 10 seconds and 60°C for 30 seconds. A melt curve analysis was performed to ensure specificity by confirming the absence of off-target amplification. Gene expression levels were quantified using the 2-ΔΔCT method, with normalization to the ACTIN2 reference gene. The list of primers is provided in Table S2.

### Transcriptomic analyses

Approximately 0.5 mm of tissue from the wound site of Arabidopsis wild type leaf petioles treated with high (0.75% agar) or low (2% agar) water conditions was collected at 0, 3, 6, 24, and 72 hours post-wounding. Samples were collected in three biological replicates, with approximately 60 petioles combined per sample. Total RNA was extracted using a Roti-Prep RNA MINI Kit according to the manufacturer’s instructions. mRNA library preparation (poly A enrichment) and sequencing (NovaSeq X Plus in 150bp paired-end) were performed at Novogene.

For RNAseq analyses, low-quality reads were filtered out using Fastp(*44*), and the cleaned reads were mapped to the Arabidopsis reference genome (TAIR10) with Hisat2 (*45*). The read counts were determined using HTSeq(*46*). Differentially expressed genes (DEGs) were identified with the DESeq2 R package (*47*), using a threshold q-value < 0.05. Genes with a q-value < 0.05 and an absolute log2(fold change) >1 were considered to exhibit statistically significant expression differences between samples. At each time point, the high water conditions compared to low water conditions. For each sample comparison at a given time point, the time 0h sample was used as the reference (high or low water conditions vs 0h). Gene expression patterns under different water conditions were analyzed using the Mfuzz package (*48*) in R. Initially, DEGs from all time points were compiled into a single list for analysis. Time 0 samples were also included to identify gene expression patterns under high and low water conditions based on the DEGs list. Gene ontology (GO) annotation was conducted using the clusterProfiler package (*49*). Enriched GO terms were identified with an adjusted p-value of less than 0.05, an enrichment fold change greater than 2, and a DEG count above 10. The lists of DEGs and GO annotations are provided in Data S1 and S2.

### Statistical Analysis

Statistical analysis was conducted using Excel version 16.77.1. A two-tailed Student’s t-test was performed between two groups within Excel. Statistical significance was determined by p-values < 0.05.

## Acknowledgements

We thank the Nottingham Arabidopsis Seed Center (NASC), Ari Pekka Mähönen (University of Helsinki, Finland), Laura Ragni (University of Freiburg, Germany), Lin Xu (CAS Center for Excellence in Molecular Plant Sciences, China), Malcolm J Bennett (University of Nottingham, UK), Peter Etchells (Durham University, UK), Peter Marhavý (Swedish University of Agricultural Sciences, Sweden), Stéphanie Robert (Swedish University of Agricultural Sciences, Sweden), Thomas Greb (University of Heidelberg, Germany) and Thomas Laux (University of Freiburg, Germany) for sharing materials. We thank Cecilia Wärdig and Frauke Augstein for technical assistance.

## Funding

A.K. was supported by the Horizon Europe Marie Sklodowska-Curie Actions framework HORIZON-MSCA-2021-PF (UMOCELF-101069157). A.K.V.W. was supported by a Vetenskapsrådet Grant (2022-03018). A.K., E.F. and C.W.M. were supported by a H2020 European Research Council Starting Grant (GRASP-805094). C.W.M. was supported by a Knut and Alice Wallenberg Stiftelse Wallenberg Academy Fellowship (2022-0193).

## Author contributions

A.K. and C.W.M. conceptualized the study. A.K., A.K.V.W., G.W. and E.F. performed experiments. A.Z. performed the bioinformatic analyses. A.K., A.K.V.W., A.Z. and G.W. analysed the data. A.K. and C.W.M. wrote the manuscript. All authors edited the manuscript.

## Competing interests

The authors declare no competing interests.

## Data and materials availability

There are no material restrictions or transfer agreements. All data are accessible in the main text and supplementary materials. Transcriptomic data reported in this paper are deposited in the NCBI Sequence Read Archive (SRA) with BioProject ID PRJNA1129885. The transcriptomic data is available from SRA. This study did not generate original code.

---

**Table S1.** List of Arabidopsis mutants and transgenic lines used in this study.

| Genotype | Background | Reference |
| --- | --- | --- |
| *arf6-2* | Columbia | Nagpal et al.(50) |
| *arf7-1* | Columbia | Okushima et al.(51) |
| *arf8-3* | Columbia | Nagpal et al.(50) |
| *arf7-1,arf19-1* | Columbia | Okushima et al (51) |
| *ein2-1* | Columbia | Guzmán and Ecker (52) |
| *eto2* | Wassilewskija | Vogel et al. (30) |
| *eto3* | Columbia | Woeste et al. (53) |
| *etr1-1* | Columbia | Bleecker et al. (29) |
| *lbd1,lbd11* | Columbia | Ye et al. (10) |
| *lbd1,lbd3,lbd4,lbd11* | Columbia | Ye et al. (10) |
| *lbd16-1* | Columbia | Okushima et al.(9) |
| *slr-1* | Columbia | Fukaki et al.(54) |
| *shy2-31* | Landsberg *erecta* | Knox et al.(55) |
| *wox4-1* | Columbia | Hirakawa et al. (56) |
| *wox14-1* | Columbia | Etchells et al. (57) |
| *wox4-1,wox14-1* | Columbia | Etchells et al. (57) |
| *wox13-2,wox14-1* | Columbia | Sakakibara et al. (58) |
| *yuc2,yuc5,yuc8,yuc9* | Columbia | (Müller-Moulé et al. (59) |
| *yuc1D* | Columbia | Weigel et al. (60) |
| *35S::XVE>>LBD11* | Columbia | Ye et al.(10) |
| *35S::LBD16* | Columbia | (Orosa-Puente et al. (3) |
| *35S::PLT2-GR* | Columbia | Galinha et al.(61) |
| *35S::WOX11* | Columbia | Liu et al.(18) |
| *35S::XVE>>WOX4* | Columbia | Zhang et al.(11) |
| *ACS6<sub>pro</sub>::NLS-3xVenus* | Columbia | Marhavý et al.(62) |
| *JAZ10<sub>pro</sub>::NLS-3xVenus* | Columbia | Marhavý et al.(62) |
| *DR5rev:3xVenus-N7* | Columbia | Heisler et al.(63) |
| *LBD1<sub>pro</sub>::erYFP* | Columbia | Ye et al.(10) |
| *LBD11<sub>pro</sub>::erYFP* | Columbia | Ye et al.(10) |
| *WOX4<sub>pro</sub>::erYFP* | Columbia | Suer et al.(64) |
| *LBD16::LBD16-GFP* | Columbia | Orosa-Puente et al.(3) |
| *PLT2::PLT2-YFP* | Columbia | Mähönen et al.(65) |
| *SHY2/IAA2<sub>pro</sub>::NLS-3x-VENUS* | Columbia | Vermeer et al.(66) |
| *WOX11<sub>pro</sub>::H2B-eGFP* | Columbia | Zhai and Xu(67) |
| *PXY<sub>pro</sub>::erECFP* | Columbia | Agusti et al. (68) |
| *YUC4<sub>pro</sub>::NLS-3xGFP* | Columbia | Robert et al.(69) |

**Table S2.**
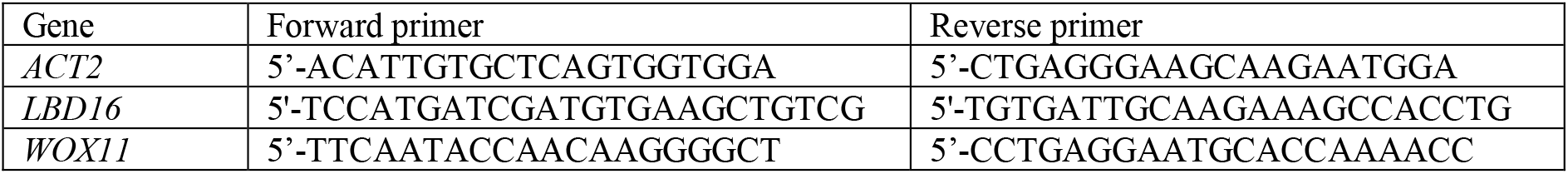
Primers Used for Quantitative Real-Time PCR.

**Data S1. (separate file)**

Differentially expressed genes (DEGs) during regeneration under varied water conditions. Analysis of differentially expressed genes (DEGs) during regeneration at different time points under high and low water conditions, compared to the 0-hour time point.

**Data S2. (separate file)**

Gene ontology (GO) annotation of gene clusters.

Gene ontology (GO) annotation analysis of various gene clusters.

**Fig. S1:**
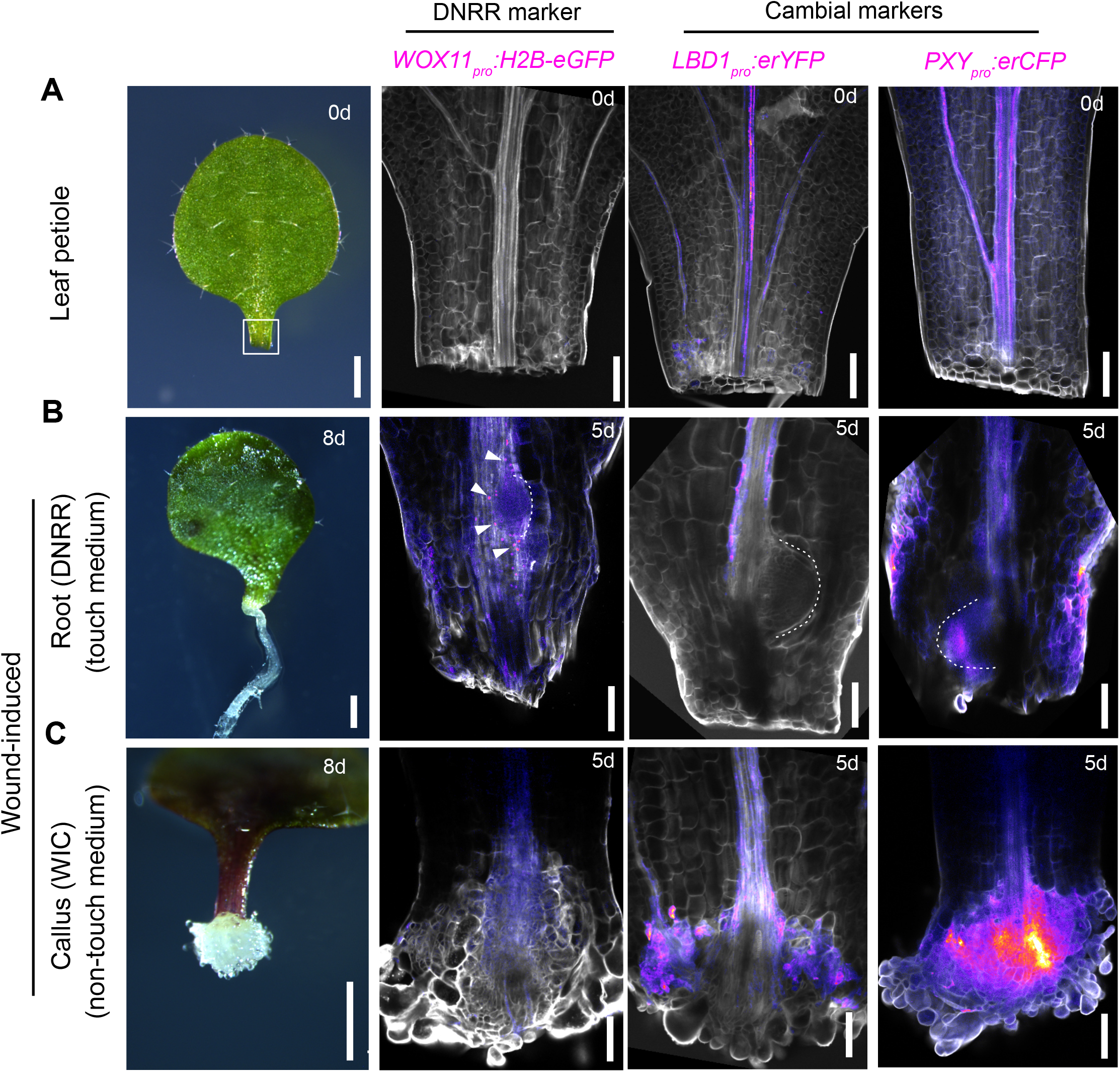
Wound-induced callus activates cambium-related markers but not root regeneration-related markers. (**A-C**) Differential expression patterns of fluorescence reporters (magenta) for *WOX11* (*de novo* root regeneration, DNRR, marker), and *LBD1* and *PXY* (cambial markers) in DNRR and wound-induced callus (WIC) at various days (d) after wounding. Calcofluor white (grey) was used to stain cell walls. Scale bar: 500µm for the bright-field images, 100µm for the confocal images.

**Fig. S2:**
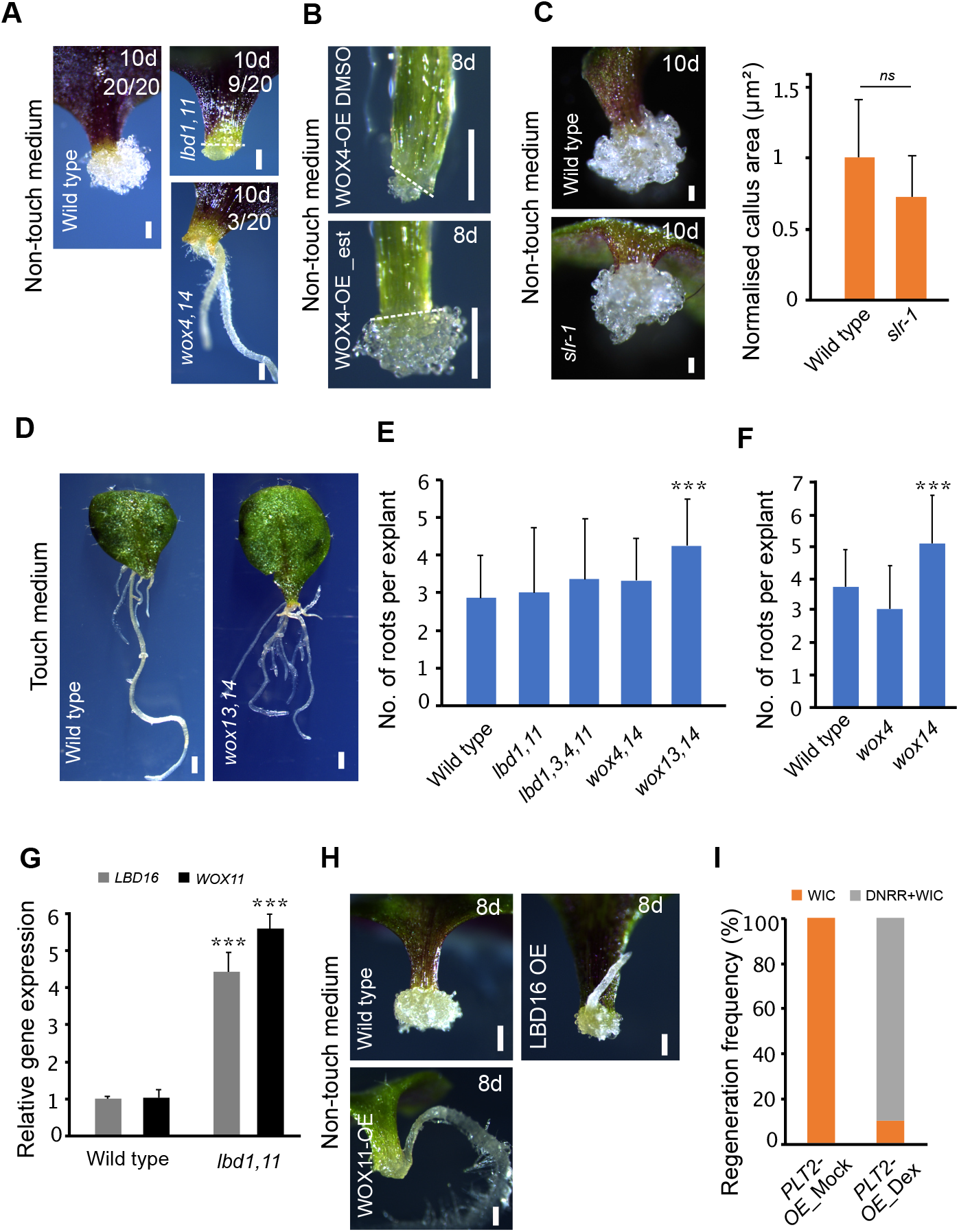
A trade-off between wound-induced callus and root regeneration determines regeneration fates. (**A**) Formation of wound-induced callus (WIC) or *de novo* root regeneration (DNRR) in leaf explants of wild type and cambial mutants with wound sites not in contact with the medium. (**B**) WIC formation in the hypocotyl explants of *WOX4-OE* with the wound site not touching the medium. (**C**) Images and quantifications of normalized WIC areas of wild type and *slr-1* cotyledon petioles with the wound site not touching the medium (n=6-8 explants per genotype; mean±SD; Student’s t-test). (**D**) Representative images showing DNRR in the *wox13,14* mutant with the wound site touching the medium. (**E**) & (**F**) Frequency of DNRR in various mutants compared to wild type with the wound site touching the medium. (n=28-31 explants in (**E**) and 21-36 explants in (**F**) per genotype; mean±SD; ***p<0.001; Student’s t-test). (**G**) *LBD16* and *WOX11* transcript levels in *lbd1,11* mutant petioles 4 days post-wounding with the wound site not touching the medium, measured by RT-qPCR. Expression normalized to ACTIN2; error bars represent SD from three independent replicates (***p<0.001; Student’s t-test). (**H**) DNRR formation in *LBD16-OE* (constitutive) and *WOX11-OE* (constitutive) with the wound site not touching the medium. (**I**) DNRR and WIC formation frequencies in the petiole of *PLT2-OE* with the wound site not touching the medium. Mock (ethanol) and 5µM dexamethasone (DEX) treatments are shown (n=10 explants per genotype; mean±SD; student’s t-test). Scale bar: 500µm in (**D**) and 200µm in the rest.

**Fig. S3.**
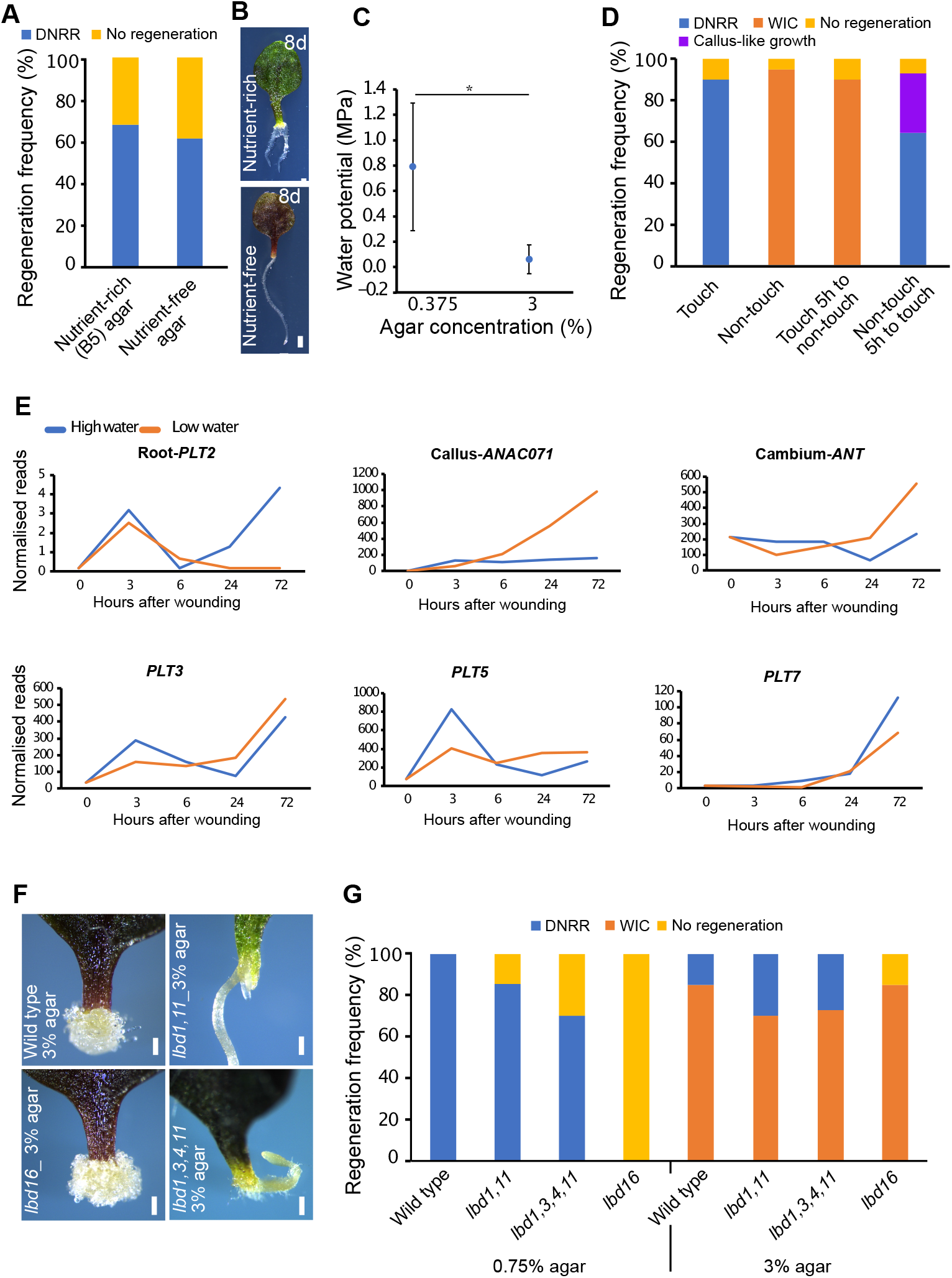
Water availability determines regeneration fates. (**A**) Regeneration frequencies in petioles with the wound site touching either nutrient rich agar medium or nutrient-free agar medium 8 days post wounding. (n=18-19 explants per treatment). (**B**) Root regeneration in petioles exposed to either nutrient-rich or nutrient-free agar media. (**C**) Correlation between water potential (MPa) and agar concentration of B5 medium. (**D**) Regeneration frequencies after changing the tissue orientation from touch to non-touch or non-touch to touch 5 hours post wounding in Arabidopsis leaf petioles. (**E**) Transcriptional dynamics of genes associated with cambium and regeneration at high (0.75% agar) and low (3% agar) water conditions. (**F**) Wound-induced callus (WIC) or *de novo* root regeneration (DNRR) in cambial and DNRR mutants under low water (3% agar) conditions. (G) Regeneration frequencies in cambial and DNRR mutants under high (0.75% agar) and low (3% agar) water conditions. (n=10-20 explants per genotype). Scale bar: 500µm in (B), 200µm in (F)

**Fig. S4.**
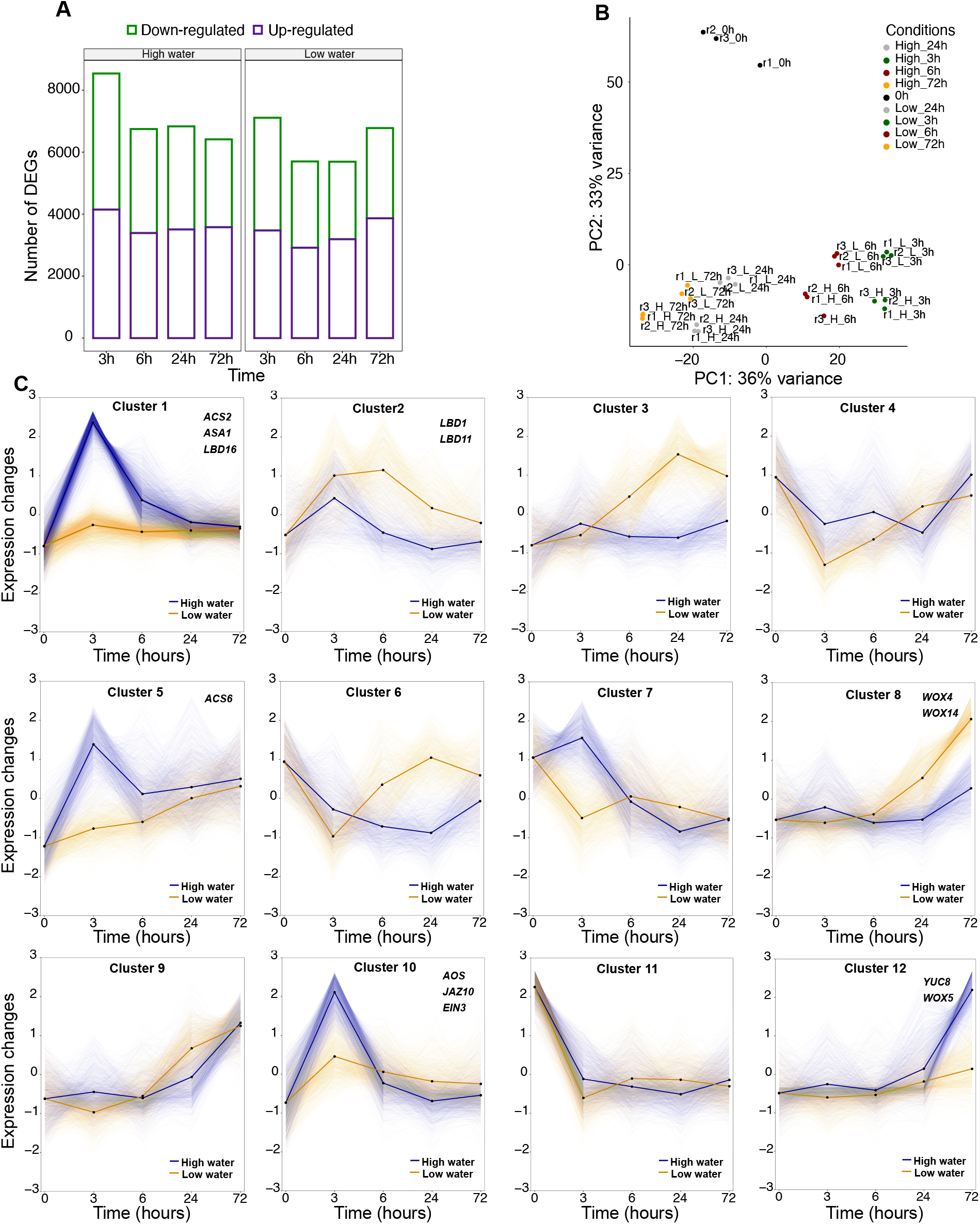
Transcriptional dynamics of regeneration fate changes. (**A**) The distribution of differentially expressed genes (DEGs) in response to high and low water conditions. (**B**) Principal Component Analysis (PCA) of the gene expression data from the regeneration fate transcriptome. The two principal components (PC1 and PC2) explain 69% of the total variation in the regeneration transcriptome. Different colors indicate various time points after wounding (in hours). Data from three biological replicates (r1, r2, r3) per water condition (high (H) or low (L) water) per time point are shown. (**C**) Clustering analysis of transcriptional dynamics under high and low water availability conditions (12 clusters in total). Lines represent the average expression levels of DEGs within each cluster. Representative genes for selected clusters are shown in the top right corner.

**Fig. S5.**
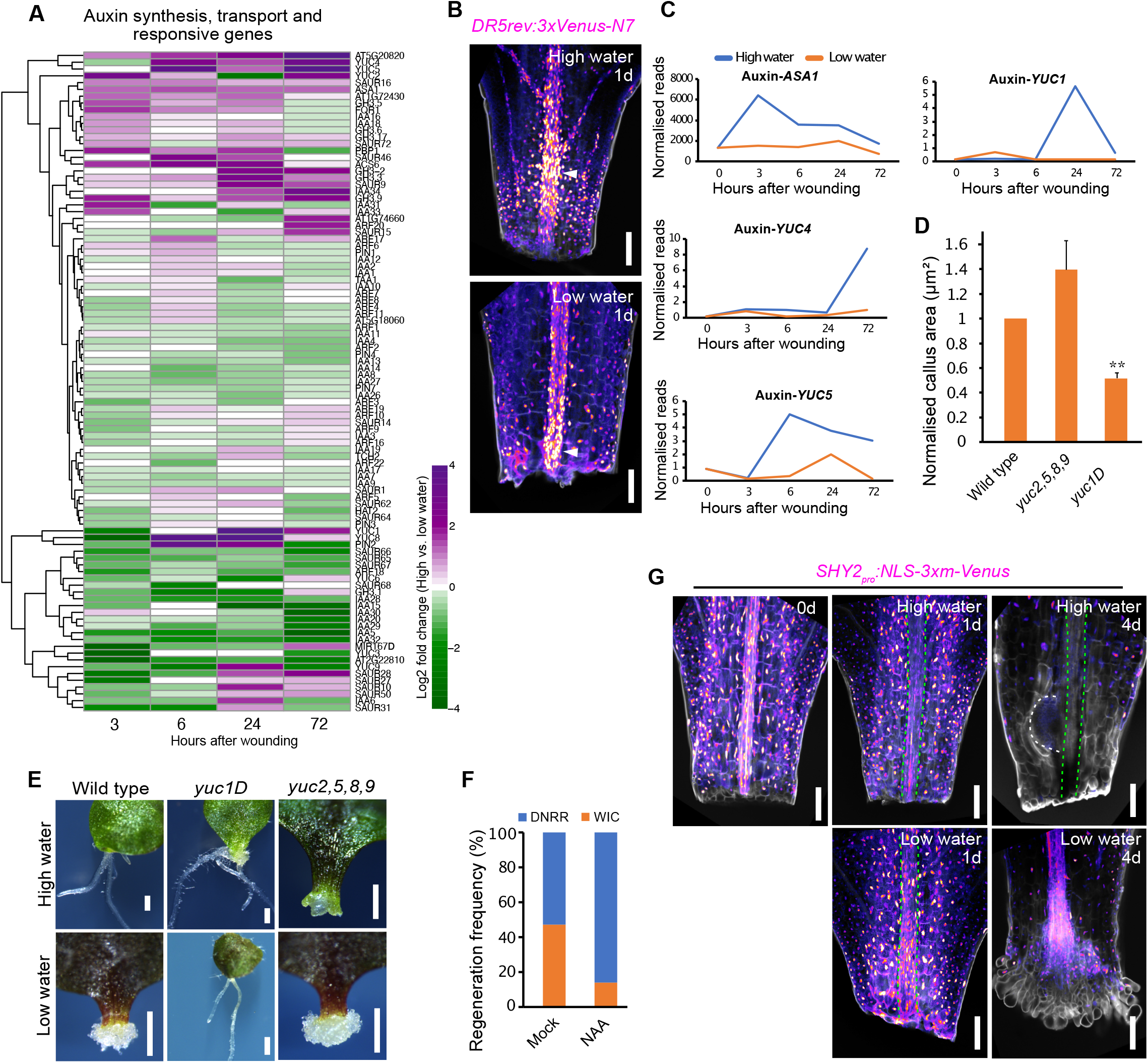
Water availability influences auxin biosynthesis and signalling during regeneration. (**A**) Heatmap showing fold changes of expression of genes involved in auxin synthesis, transport and response under high (0.75% agar) vs. low (2% agar) water conditions in wounded Arabidopsis petioles. (**B**) Expression of *DR5-VENUS* under high (0.75% agar) and low (2% agar) water conditions 1d post wounding. (**C**) Transcriptional dynamics of auxin biosynthesis genes at high and low water conditions. (**D**) Normalised wound-induced callus (WIC) area in *yuc1D* and *yuc2,5,8,9*. (n=15-20 explants per genotype; mean±SD; **p<0.01; Student’s t-test). (**E**) WIC and *de novo* root regeneration in *yuc1D* and *yuc2,5,8,9* mutants at high (0.75% agar) and low (2% agar) water conditions. (**F**) Regeneration frequencies after 0.2µM NAA treatment under intermediate water levels (1.5% agar). (n=14-19 explants per treatment). (**G**) Spatial expression of *SHY2-VENUS* under high (0.75% agar) and low (2% agar) water conditions. Green dotted lines mark vasculature, while the white dotted line marks the root primordium. Scale bar: 100µm in (B and G) and 500µm in (E).

**Fig. S6.**
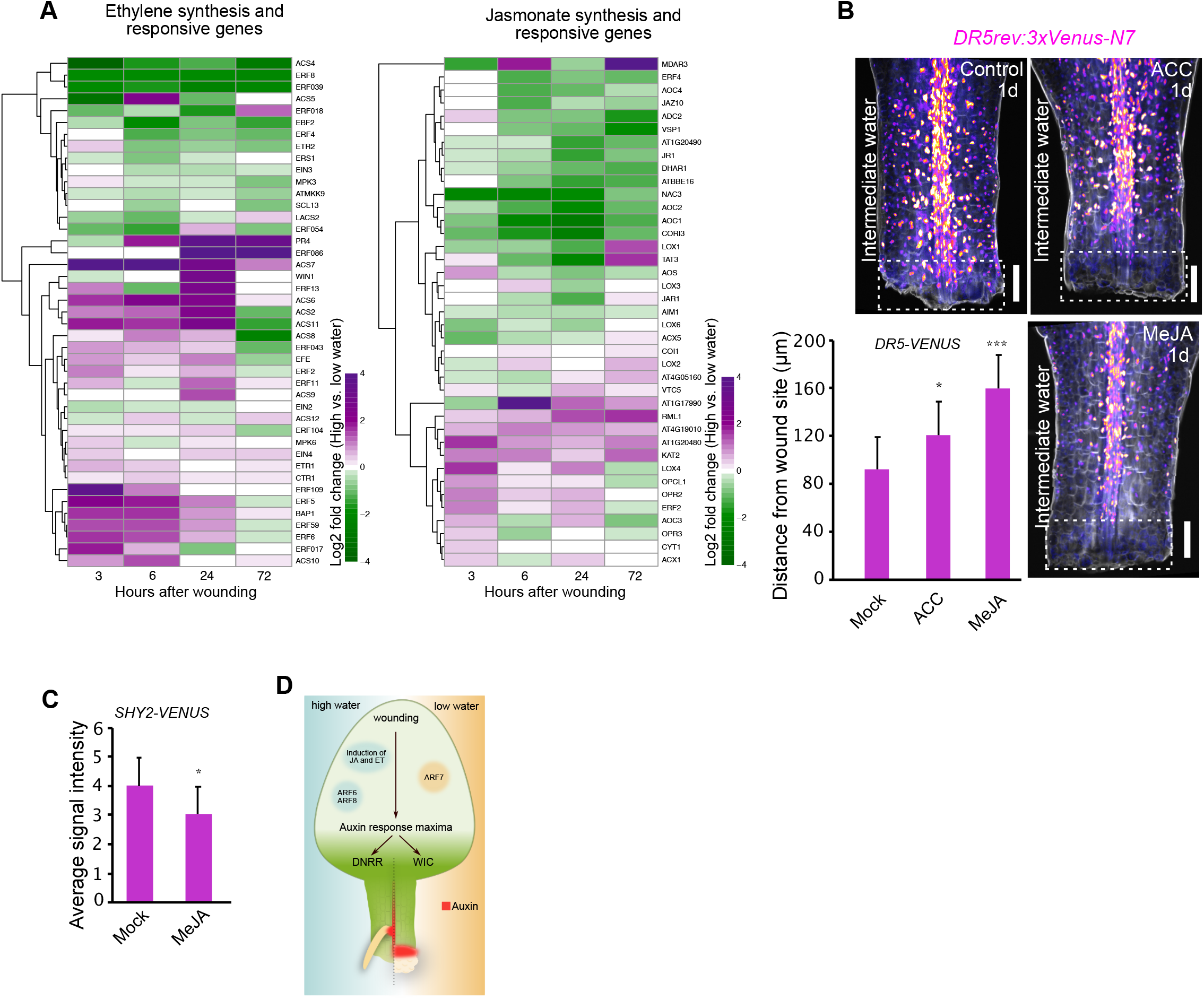
Ethylene and jasmonic acid influence spatial auxin distribution. (**A**) Heatmaps showing fold changes of expression of genes involved in ethylene, and jasmonate synthesis and response under high (0.75% agar) vs. low (2% agar) water conditions in wounded Arabidopsis petioles. (**B**) *DR5-VENUS* shows auxin response changes (denoted in white box) after ethylene (4µM ACC) or methyl-jasmonate (5µM MeJA) treatment at intermediate water conditions (1.5% agar). Auxin response is also quantified at the wound site after ACC and MeJA treatment for 24h at intermediate water availability (1.5% agar), measured by the distance to the visible high auxin response. (n=9-10 explants per treatment; mean±SD; *p<0.05; ***p<0.001; Student’s t-test). (**C**) Average signal intensity of *SHY2-VENUS* post-treatment with 5µM MeJA or mock for 43h at intermediate water levels (1.5% agar). (n=9-10 explants per treatment; mean±SD; *p<0.05; student’s t-test). (**D**) Working model showing water availability determines regeneration fates in detached leaf tissues. High water triggers *de novo* root regeneration (DNRR), while low water promotes wound-induced callus (WIC) formation by modulating the spatial distribution of auxin response maxima, regulated by water-induced ethylene and jasmonate responses. Scale bar: 100µm.

## Notes

### Competing Interest Statement

The authors have declared no competing interest.

